# PRH1 mediates ARF7-LBD dependent auxin signaling to regulate lateral root development in *Arabidopsis thaliana*

**DOI:** 10.1101/558387

**Authors:** Feng Zhang, Wenqing Tao, Ruiqi Sun, Junxia Wang, Cuiling Li, Xiangpei Kong, Huiyu Tian, Zhaojun Ding

## Abstract

The development of lateral roots in *Arabidopsis thaliana* is strongly dependent on signaling directed by the AUXIN RESPONSE FACTOR7 (ARF7), which in turn activates LATERAL ORGAN BOUNDARIES DOMAIN (LBD) transcription factors (*LBD16*, *18*, *29* and *33*). Here, the product of *PRH1,* a *PR-1* homolog annotated previously as encoding a pathogen-responsive protein, was identified as a target of ARF7-mediated auxin signaling and also as participating in the development of lateral roots. *PRH1* was shown to be strongly induced by auxin treatment, and plants lacking a functional copy of *PRH1* formed fewer lateral roots. The transcription of *PRH1* was controlled by the binding of both ARF7 and LBDs to its promoter region. An interaction was detected between PRH1 and GATA23, a protein which regulates cell identity in lateral root founder cells.

**Author Summary:** In *Arabidopsis thaliana* AUXIN RESPONSE FACTOR7 (ARF7)-mediated auxin signaling plays a key role in lateral roots (LRs) development. The *LATERAL ORGAN BOUNDARIES DOMAIN (LBD)* transcription factors (*LBD16*, *18*, *29* and *33*) act downstream of ARF7-mediated auxin signaling to control LRs formation. Here, the *PR-1* homolog *PRH1* was identified as a novel target of both ARF7 and LBDs (especially the LBD29) during auxin induced LRs formation, as both ARF7 and LBDs were able to bind to the *PRH1* promoter. More interestingly, PRH1 has a physical interaction with GATA23, which has been also reported to be up-regulated by auxin and influences LR formation through its regulation of LR founder cell identity. Whether the interaction between GATA23 and PRH1 affects the stability and/or the activity of either (or both) of these proteins remains an issue to be explored. This study provides improves new insights about how auxin regulates lateral root development.

## Introduction

The architecture of the root system depends on the density of lateral roots (LRs) formed along with the extent of root branching. LRs are initiated from mature pericycle cells lying adjacent to the xylem pole, referred to in *Arabidopsis thaliana* as the xylem pole pericycle [1, 2]. A subset of these cells, namely the founder cells, undergo a series of highly organized divisions to form an LR primordium, which eventually develops into an LR. The entire process of LR development, from its initiation to its emergence, is regulated by auxin [1, 3, 4]. The accumulation of auxin in protoxylem cells results in the priming of neighboring pericycle cells to gain founder cell identity, forming an LR pre-branch site [5, 6]. This auxin signaling is mediated by the proteins ARF7 and ARF19, which act in the IAA28-dependent pathway [7, 8]. LR initiation depends on the transcriptional activation, through the involvement of the transcription factor *LBD16* and *LBD18*, of either *E2Fa* or *CDK A1;1* and *CYC B1;1* [9, 10]. Other downstream targets such as EXP14 and EXP17 and the products of certain cell wall loosening-related genes are also either directly or indirectly regulated by LBD18 to promote the emergence of an LR [10, 11, 12]. LBD18 has been also found to control LR formation through its interaction with GIP1, which results in the transcriptional activation of *EXP14* [13].

LBD transcription factors are among the most well studied downstream targets of ARF7 and ARF19 [14]. ARF7 has also been found to control LR emergence through its regulation of *IDA* and genes which encode leucine-rich repeat receptor-like kinases such as *HAE* and *HSL2* [15]. These interactions take place in the overlaying tissues, thereby influencing cell wall remodeling and cell wall degradation during LR emergence [15]. The product of *PRH1* (*At2g19990*), a homolog of *PR-1*, is currently annotated as encoding a pathogen responsive protein [16]. Here, this protein has been identified as a novel target of ARF7-mediated auxin signaling, and evidence is provided of its involvement in LR development.

## Results

### PRH1 is a target of ARF7-mediated auxin signaling

To identify additional targets involved in ARF7-mediated auxin signaling and LR development (Fig 1A and 1B), a comparison was made between the root transcriptomes of eight-day-old wild type (WT) or *arf7* mutant seedlings exposed to auxin treatment respectively. A total of 74 genes was classified as DEGs (P < 0.0001, log_2_ fold change >1) in WT but not in *arf7* mutant, and these were then compared to the *arf7* and WT transcriptome after auxin treatment (Fig 1C and S1A Fig). Twenty-three DEGs (P < 0.0001, log_2_ fold change >1.5) were selected as the candidate target genes induced by auxin, which were also dependent on ARF7 (S1B Fig). We then compared the expression patterns of these 23 candidate genes in lateral root initiation using Arabidopsis eFP Browser (http://bar.utoronto.ca/efp/cgi-bin/efpWeb.cgi). As a result of this analysis, eight genes were found to be involved in LR development. The most strongly differentially transcribed of these genes was *PRH1* (*At2g19990*) (S1C Fig).

**Fig 1.**
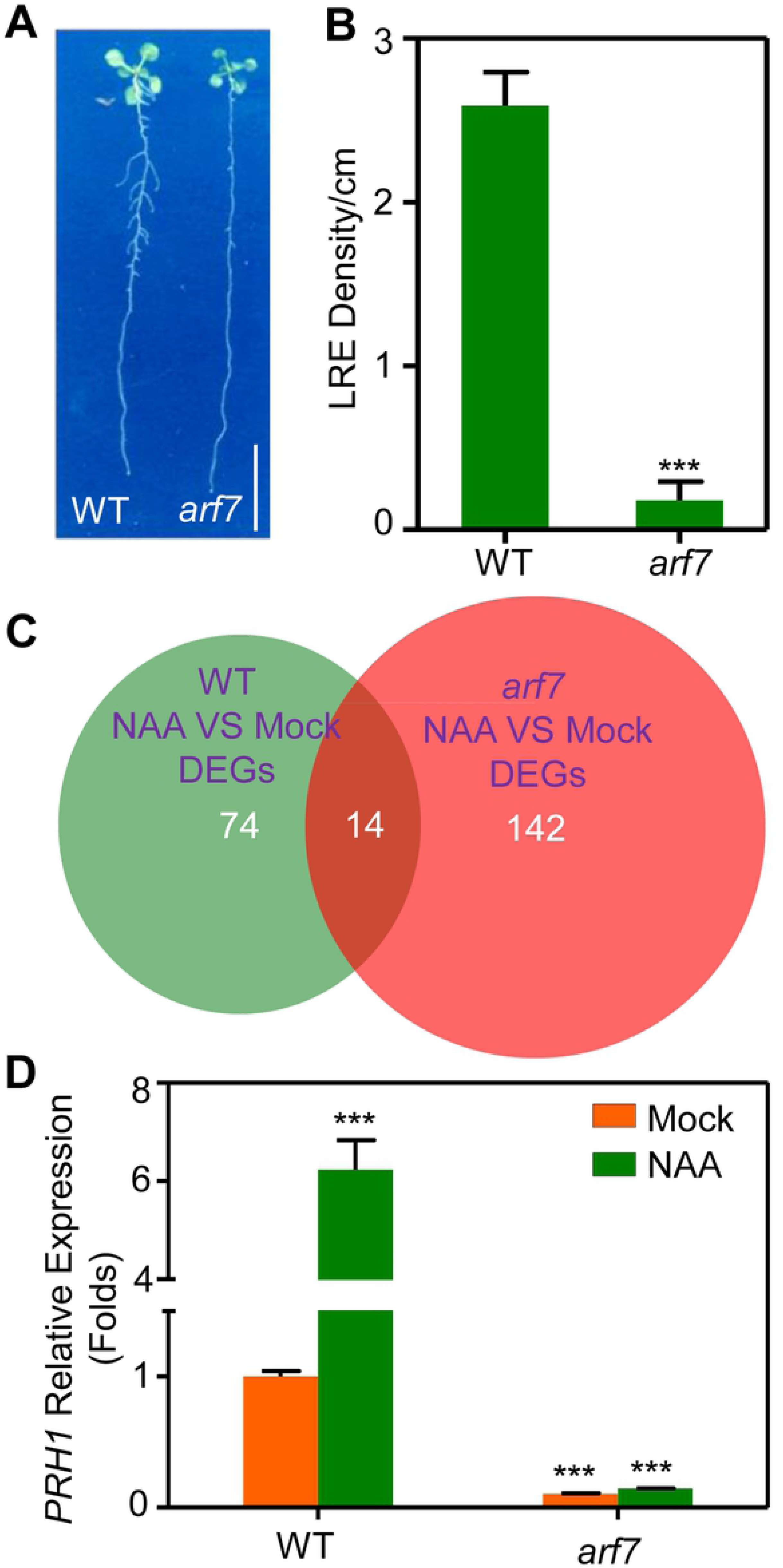
*PRH1* is a downstream target of ARF7-mediated auxin signaling. (A) Eight-day-old seedlings of WT and *arf7*. Scale bar: 1 cm. (B) The ratio between the number of LRs and the length of the primary root of WT and *arf7* seedlings. Data shown as mean±SE, three biological replicates in the experiment, 20 plant seedlings for each repeat. ***: the mutant’s performance differs significantly (*P*<0.001) from that of WT. (C) RNA-seq analysis: RNA was extracted from the roots of eight-day-old seedlings exposed or not exposed to auxin. DEGs: differentially expressed genes, NAA: naphthalene acetic acid. (D) Transcription level of *PRH1* in the roots of eight-day-old WT and *arf7* seedlings exposed (green) or not exposed (orange) to 10 μmol NAA for 4 hours. Transcript abundances assayed by qPCR. Values represent averages of three biological replicates in the experiment, and the total RNA was extracted from about 100 seedlings for each repeat. Error bars represent SE, ***: the mutant’s performance differs significantly (*P*<0.001) from that of WT.

*PRH1*, a homolog of *PR-1* which has been annotated as encoding a pathogen-responsive protein [16]. Though PRH1 was highly induced by the auxin treatment in WT, it was completely non-responsive to auxin treatment in *arf7* according to our qPCR assays (Fig 1D). The *GUS* expression analysis in 8-day-old seedlings carrying a *PRH1pro::GUS* transgene implied that *PRH1* is mainly expressed in the site of LR initiated, which was enhanced by exogenous auxin treatment (S2 Fig). The GUS signal was detected only at the LR primordia in WT background (Fig 2A), and the level of GUS activity was greatly enhanced following exposure of the plants to auxin (Fig 2B). When the *PRH1pro::GUS* transgene was expressed in the *arf7* mutant background, the *PRH1pro::GUS* expression was strongly suppressed, and auxin treatment only slightly induced the expression of *PRH1pro::GUS* (Fig 2C and 2D). The conclusion was that the auxin-responsive gene *PRH1* is expressed in the LR primordia and its expression is dependent on ARF7-mediated auxin signaling.

**Fig 2.**
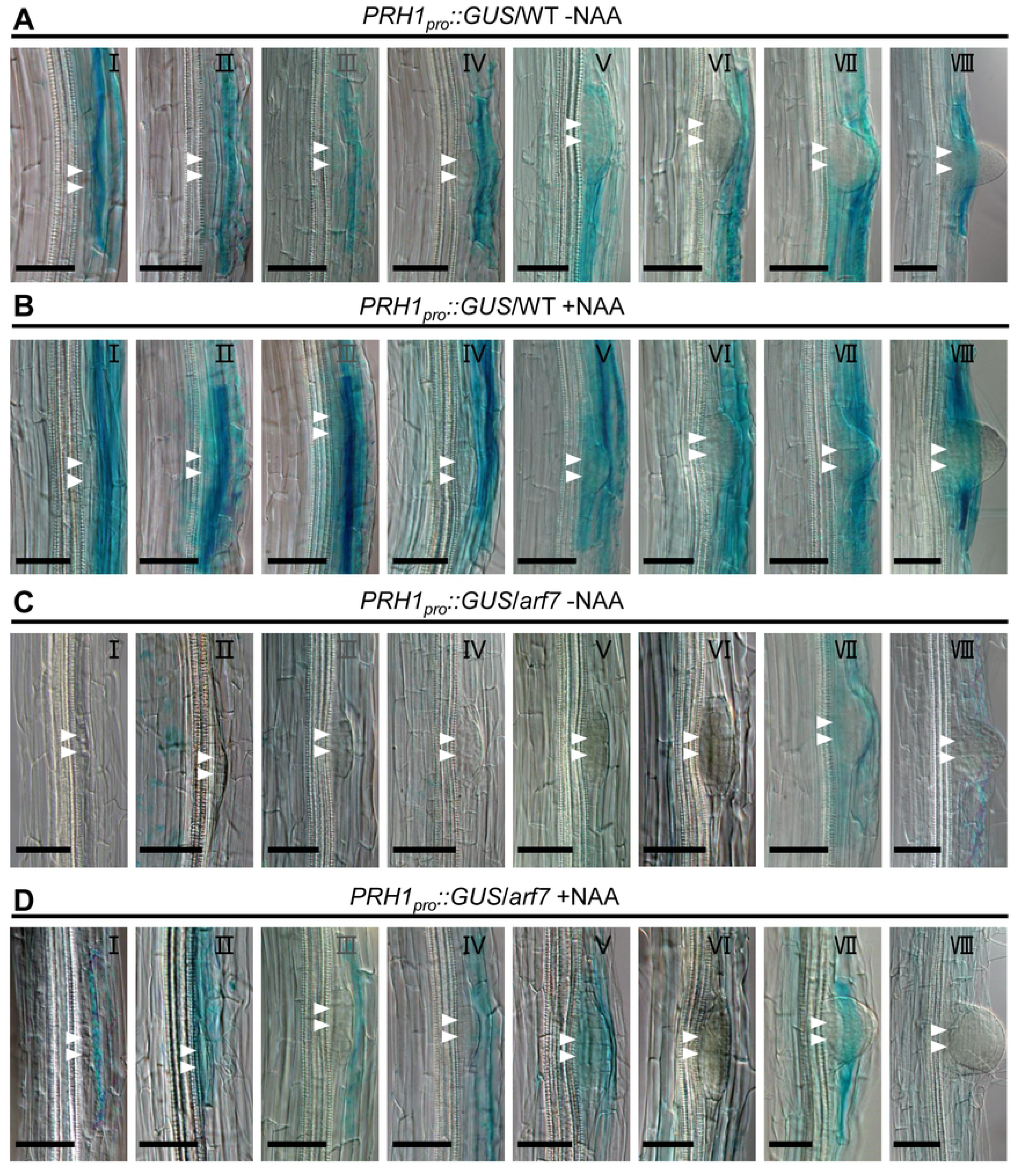
Histological localization and expression of *PRH1pro::GUS* transgene during LR formation. (A) and (B) *PRH1* is expressed in the LR primordia in WT seedlings not exposed to auxin(A), and the intensity of its expression is strongly enhanced when the plants were treated with auxin (B). (C) and (D) GUS activity is suppressed in non-treated *arf7* seedlings (C), and even more so in treated ones (D). NAA: naphthalene acetic acid. Before GUS staining the seedlings were soaked for 4 hours in liquid half strength Murashige and Skoog (MS) medium, which containing 10 μmol NAA or not. The developmental stages (I through VIII) of the LRs are indicated in the bottom left corner. GUS (β-glucuronidase) signals are shown in blue. Scale bars: 100 μm.

### PRH1 acts downstream of ARF7 to regulate LR development

To address if auxin-induced *PRH1* expression is involved into LR development, we examined the LR phenotypes in both *prh1* mutant and the PRH1 over-expression lines (S3A-3C Fig). The expression level of *PRH1* in its T-DNA insertion mutants and the over-expression lines were verified (S3D Fig). Examination of LR development in *prh1-1* mutant plant showed that the absence of a functional copy of *PRH1* inhibited the formation of both emerged LRs and total LRs (Fig 3A and 3B). Though LR development is not greatly affected by the over-expression of *PRH1* in a WT background, over-expression of *PRH1* could significantly increase the LR numbers of *arf7* mutant plants (Fig 3A and 3B). In summary, PRH1 acts downstream of ARF7 to regulate LR development.

**Fig 3.**
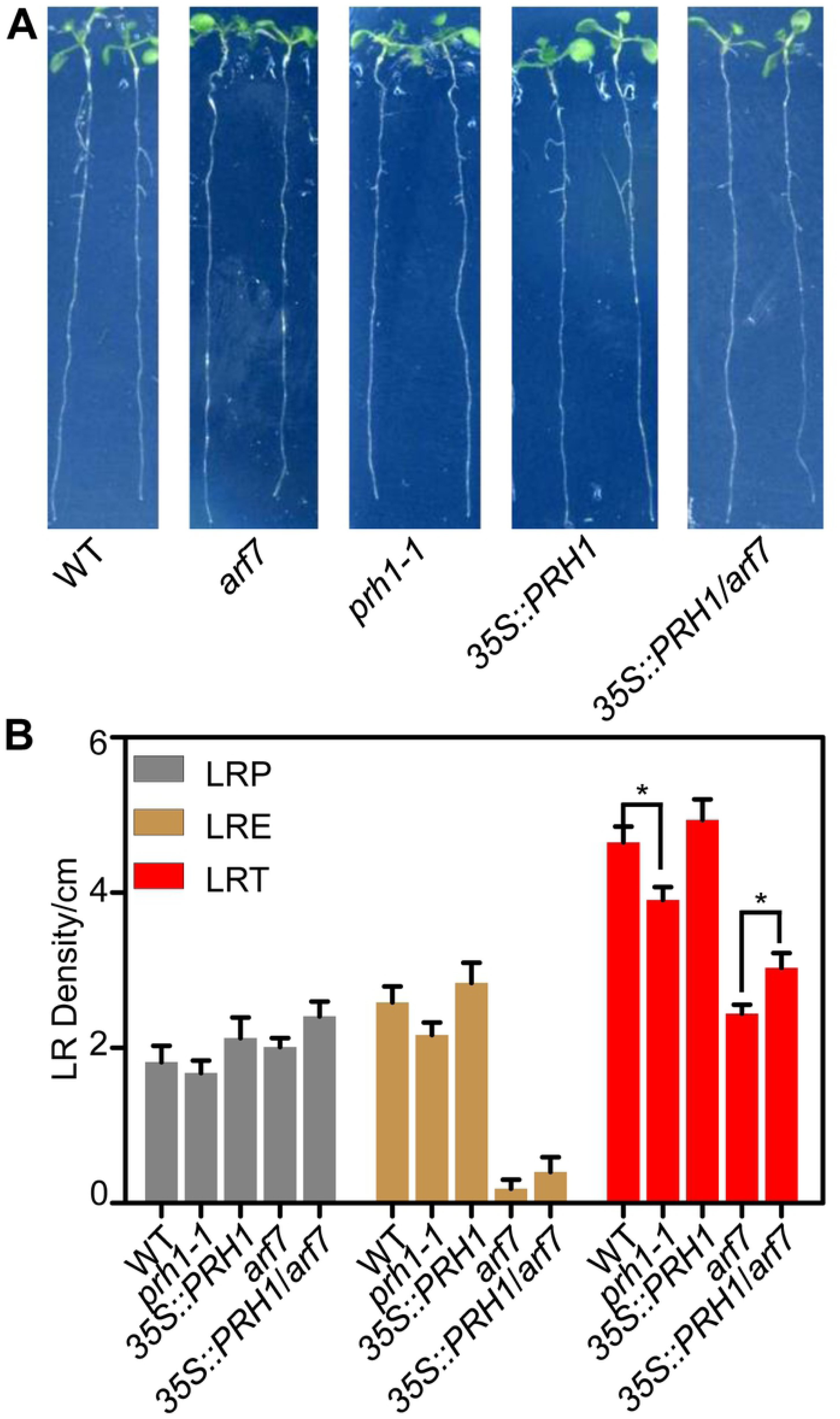
*PRH1* acts downstream of ARF7-mediated LR development. (A) Eight-day-old seedlings of WT, *arf7*, *prh1-1* mutants and transgenic lines harboring either *35S::PRH1* or *35S::PRH1/arf7.* Bar: 0.5 cm. (B) The ratio between the number of LRs and the length of the primary root. LRP: LR primordia. LRE: emerged LRs, LRT: all the LRs including the LRP and LRE. Data shown as mean±SE, three biological replicates in the experiment, 20 plant seedlings for each repeat. The sterisks indicate means which differ significantly (*P*<0.05) from one another.

### ARF7 directly regulates *PRH1* transcription

The observation that the abundance of *PRH1* transcript was reduced in the *arf7* mutant by auxin treatment (Fig 1D) suggested that the gene was transcriptionally regulated by the auxin response factor ARF7. The gene’s promoter sequence harbors the known auxin response elements (AuxREs) TGTC (Fig 4A). A transient expression assay carried out in *A. thaliana* leaf protoplasts showed that ARF7 was able to activate a construct comprising the *PRH1* promoter fused with *LUC*. The co-expression of *PRH1pro::LUC* and *35S::ARF7* had a large positive effect on the intensity of the luminescence signal (Fig 4B), consistent with the idea that ARF7 is able to activate the *PRH1pro::LUC* construct. A ChIP-qPCR assay confirmed the association of ARF7 with the AuxRE elements present on the *PRH1* promoter (Fig 4A and 4C), and a yeast one hybrid assay showed that ARF7 was able to bind to the *PRH1* promoter (Fig 4D). The conclusion was that the effect of ARF7 binding to the *PRH1* promoter was regulating the gene’s transcription, thereby influencing LR development.

**Fig 4.**
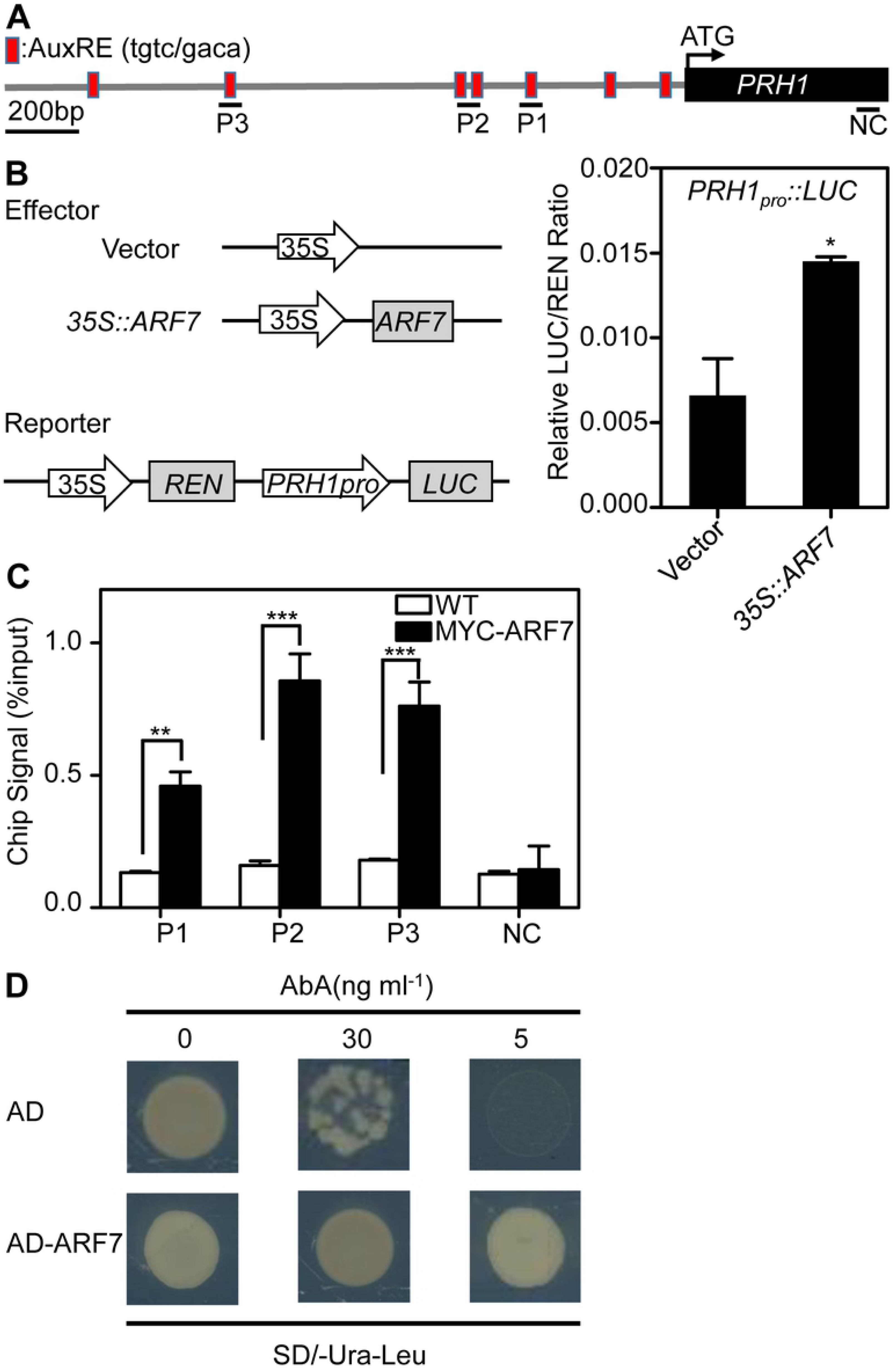
*PRH1* is regulated by *ARF7* at the transcriptional level. (A) Structure of *PRH1* promoter and the fragments used in the CHIP-qPCR assay. AuxREs are indicated by red squares, and black lines show the promoter regions containing the AuxREs used in this assay. NC: negative control. (B) ARF7 transactivates the *PRH1* promoter in *A. thaliana* leaf protoplasts. The left hand panel is a schematic of the effector (*35S::ARF7*) and reporter (*PRH1pro::LUC*) constructs. The empty vector pBI221 was used as a negative control; the right hand panel shows the ratio of *ARF7* drived *LUC* and the empty vector (negative control) to 35S promoter drived *REN* respectively. LUC: firefly luciferase activity, REN: renilla luciferase activity. Values shown as mean±SE, three biological replicates in the experiment. *: means differ significantly (*P*<0.05) from the negative control. (C) ARF7 is associated with the *PRH1* promoter according to a CHIP-qPCR assay. Chromatin isolated from a plant harboring *35S::MYC-ARF7* and a WT mock control was immunoprecipitated with anti-MYC antibody followed the amplification of regions P1, P2 and P3. The coding region segment NC was used as the negative control. The ChIP signal represents the ratio of bound promoter fragments (P1-P3) to total input after immuno-precipitation. Values shown as mean±SE, three biological replicates in the experiment. **, ***: means differ significantly (*P*<0.01, *P<*0.001) from the WT control. (D) Physical interaction of ARF7 with the *PRH1* promoter according to a Y1H assay. The plasmid pGADT7-ARF7 was introduced into Y1H Gold cells harboring the reporter gene *PRH1pro::AbAr* and the cells were grown on SD/-Ura-Leu medium in the presence of 30 or 50 ng/mL aureobasidin A (AbA). The empty vector pGADT7 (AD) was used as a negative control.

### Auxin-induced *PRH1* expression is regulated by LBD29

Members of the LBD family are known to function downstream of ARF7 to control LR initiation and emergence [14, 17, 18, 19]. When the transcription level of *PRH1* was investigated in the three single *lbd* mutants *lbd16*(Salk_040739), *lbd18* (Salk_038125) and *lbd29* (Salk_071133) combined with or without exogenous auxin treatment, the observation was that the gene was strongly repressed in the single mutants. Though the auxin induced *PRH1* expression was normal in *lbd16* or *lbd18*, the induction was highly reduced in *lbd29*, indicating LBD29 plays a crucial role in auxin induced *PRH1* expression (S4A Fig). Consistently, the over-expression of *LBD29* strongly up-regulates *PRH1* expression (S4B Fig). In addition, we also observed the up-regulation of *PRH1* expression in LBD16 or LBD18 over-expression lines (S4B Fig) These results suggest that LBDs regulate the transcription of *PRH1*, auxin induced expression of *RPH1* is highly dependent on LBD29.

### LBDs regulate *PRH1* by binding to its promoter

The transcriptional response of *PRH1* to the loss-of-function and over-expression of certain LBDs implied that these transcription factors can also regulate *PRH1*. Though the *PRH1* promoter has the typical LBD protein binding elements including CGGCG and CACGTG (S5A Fig). Although the yeast one hybrid assay produced no direct evidence for the binding of LBD proteins to the *PRH1* promoter (S5B Fig), but the ChIP-qPCR assay did support the association of LBD16, LBD18 and LBD29 with the *PRH1* promoter (Fig 5A). Thus a transient expression assay in *A. thaliana* leaf protoplasts was used to investigate whether LBD16, LBD18 and/or LBD29 was able to influence the expression of the *PRH1pro::LUC* transgene. The outcome of co-expressing this construct with each of *35S::LBD16*, *35S::LBD18* or *35S::LBD29* was to enhance the expression of *PRH1pro::LUC* (Fig 5B). When a pair of effector transgenes (either *35S::LBD16* plus *35S::LBD18, 35S::LBD18* plus *35S::LBD29* or *35S::LBD16* plus *35S::LBD29)* was introduced, the *PRH1pro::LUC* expression was even more strongly activated (Fig 5B). The overall conclusion was that the LBDs regulate *PRH1* expression by directly binding to its promoter.

**Fig 5.**
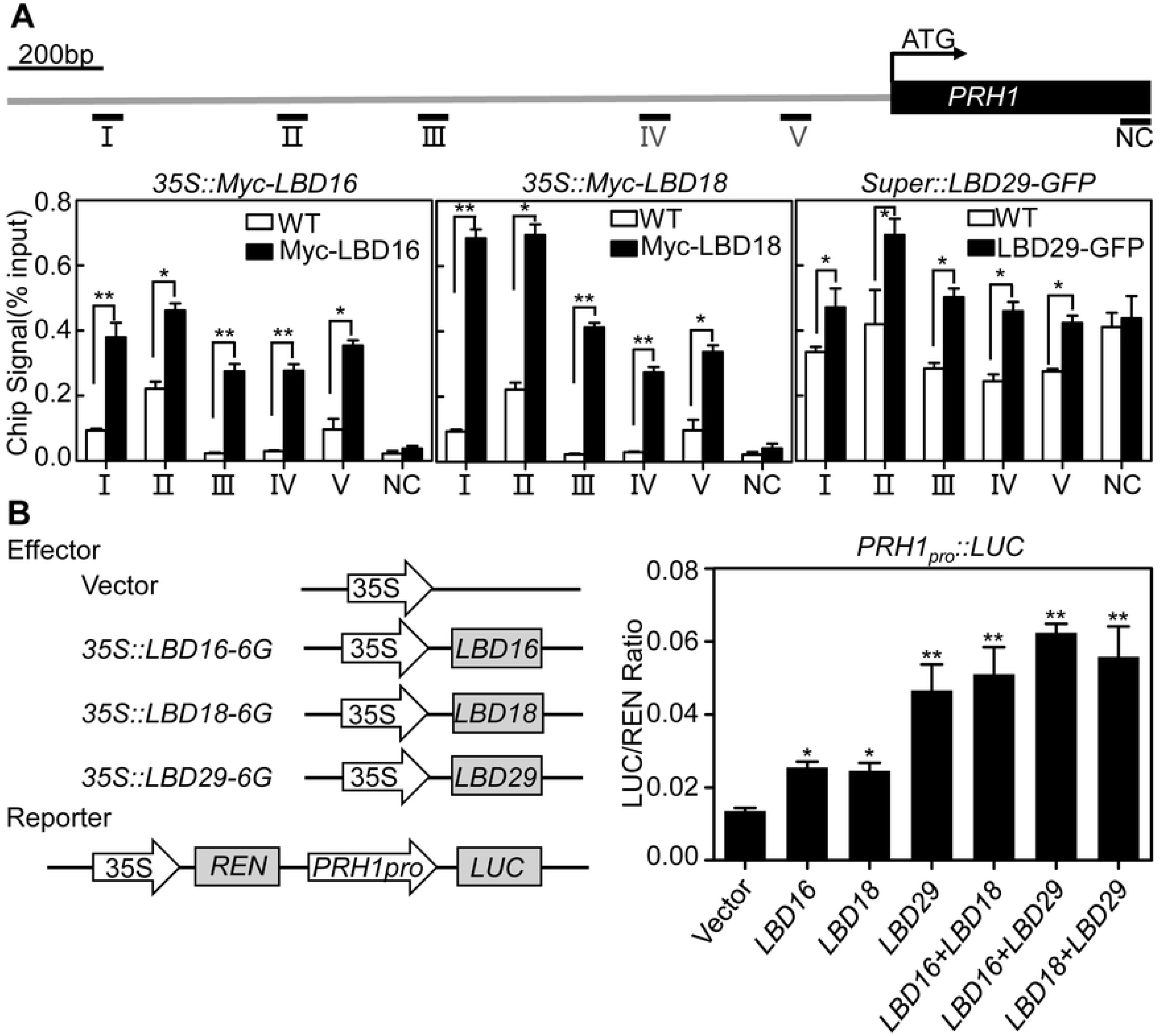
The transcription of *PRH1* is activated by LBD binding to its promoter. (A) LBDs associated with the promoter of *PRH1* according to a CHIP-qPCR assay. The upper panel shows the structure of the *PRH1* promoter, and five uniformly distributed sites (black lines) were selected for the CHIP-qPCR assay. Sites II and III include the LBD29 and LBD18 binding motif (See S5 Fig). NC: negative control. The lower panel demonstrates the ratio of bound promoter fragments (**I**-**V**;) to total input, derived from experiments involving WT seedlings (negative control) and those harboring one of the transgenes *35S::Myc-LBD16* (Myc-LBD16), *35S::Myc-LBD18* (Myc-LBD18), *Super::LBD29-GFP* (LBD29-GFP). Values shown as mean±SE, three biological replicates in the experiment. *, **: means differ significantly (*P*<0.05, *P<*0.01) from the WT control. (B) LBDs transactivate the *PRH1* promoter in *A. thaliana* leaf protoplasts as shown by a transient dual-luciferase assay. The reporter gene (*PRH1* promoter driving *LUC*) was co-transformed with one or two of the constructs *35S::LBD16*, *35S::LBD18* and *35S::LBD29.* The empty vector pBI221 was used as negative control. Values shown as mean±SD, three biological replicates in the experiment. *, **: means differ significantly (*P*<0.05, *P<*0.01) from the WT control.

### LBD regulated LR development is partially dependent on PRH1

To further study if LBDs regulated *PRH1* transcription is involved into LBD-mediated LR development, we crossed *35S::PRH1* with *lbd16*, *lbd18* or *lbd29* mutants and analyzed LR phenotypes of the generated double mutants. Compared to *lbd16*, *lbd18*, *lbd29* single mutant plants, the generated *35S::PRH1/lbd16*, *35S::PRH1 lbd18* or *35S::PRH1/lbd29* double mutants have significantly increased LR numbers (Fig 6), indicating that *PRH1* is involved into LBD-mediated LR development.

**Fig 6.**
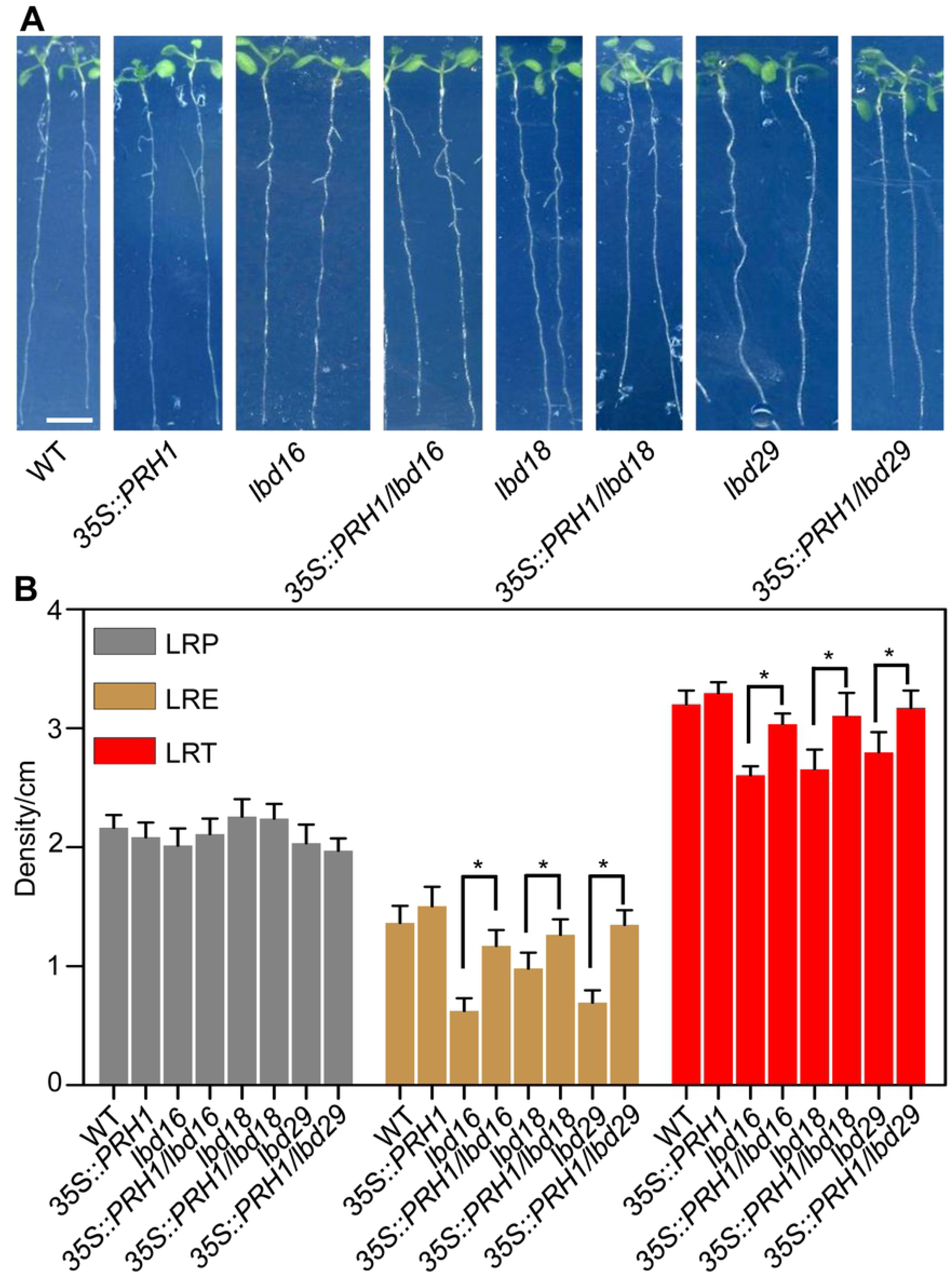
The over-expression of *PRH1* partially alleviates the defective development of LRs in the *lbd* mutant. (A) Eight-day-old seedlings of WT, the mutants *lbd16*, *lbd18* and *lbd29,* and the transgenic lines harboring *35S::PRH1*. Scale bar: 5 mm. (B) Density of LRP (LR primordia), LRE (emerged LRs) and LRT (Total LRs containing LRP and LRE) formed in eight-day-old seedlings. Data shown as means±SE, three biological replicates in the experiment, 20 plant seedlings for each repeat. *: means differ significantly (*P*<0.05) as a result of over-expressing *PRH1* in the mutant backgrounds.

### PRH1 interacts with GATA23

The subcellular localization of PRH1 is in the nuclei of *N. benthamiana* leaf cells (S6 Fig). To explore the candidate genes interacting with PRH1 in the process of regulating LR development, a yeast two hybrid screen of an *A. thaliana* root cDNA library was carried out, using *PRH1* cDNA as the bait. One of the positive clones isolated was found to include the coding sequence of *GATA23*, a gene which encodes a transcription factor known to control LR founder cell identity (Fig 7A) [8]. The interaction between PRH1 and GATA23 was confirmed using a Co-IP assay, in which C-terminal Myc-tagged full length *PRH1* cDNA (*PRH1-Myc*) and C-terminal GFP-tagged full length *GATA23* cDNA (*GATA23-GFP*) were transiently co-expressed in *A. thaliana* leaf protoplasts (Fig 7B). When the immunoprecipitated proteins were probed with an anti-GFP antibody, it was found that PRH1-Myc co-immunoprecipitated with GATA23-GFP. The suggestion was therefore that PRH1 interacts with GATA23 in planta. Further evidence for the interaction between PRH1 and GATA23 was obtained using a BiFC assay, in which a strong YFP signal was observed in the nuclei of *N. benthamiana* epidermal cells co-expressing *PRH1* fused to the N-terminal half of *YFP* and *GATA23* fused to its C-terminal half (Fig 7C). The interpretation of this result was that the auxin inducible protein PRH1 interacts with GATA23. Meanwhile the transcriptional level of GATA23 was significantly down-regulated in *arf7*, *lbd16*, *lbd18* and *lbd29* single mutant (S7 Fig). Together, auxin induced PRH1 interacts with GATA23, which has been also reported to be up-regulated by auxin [20].

**Fig 7.**
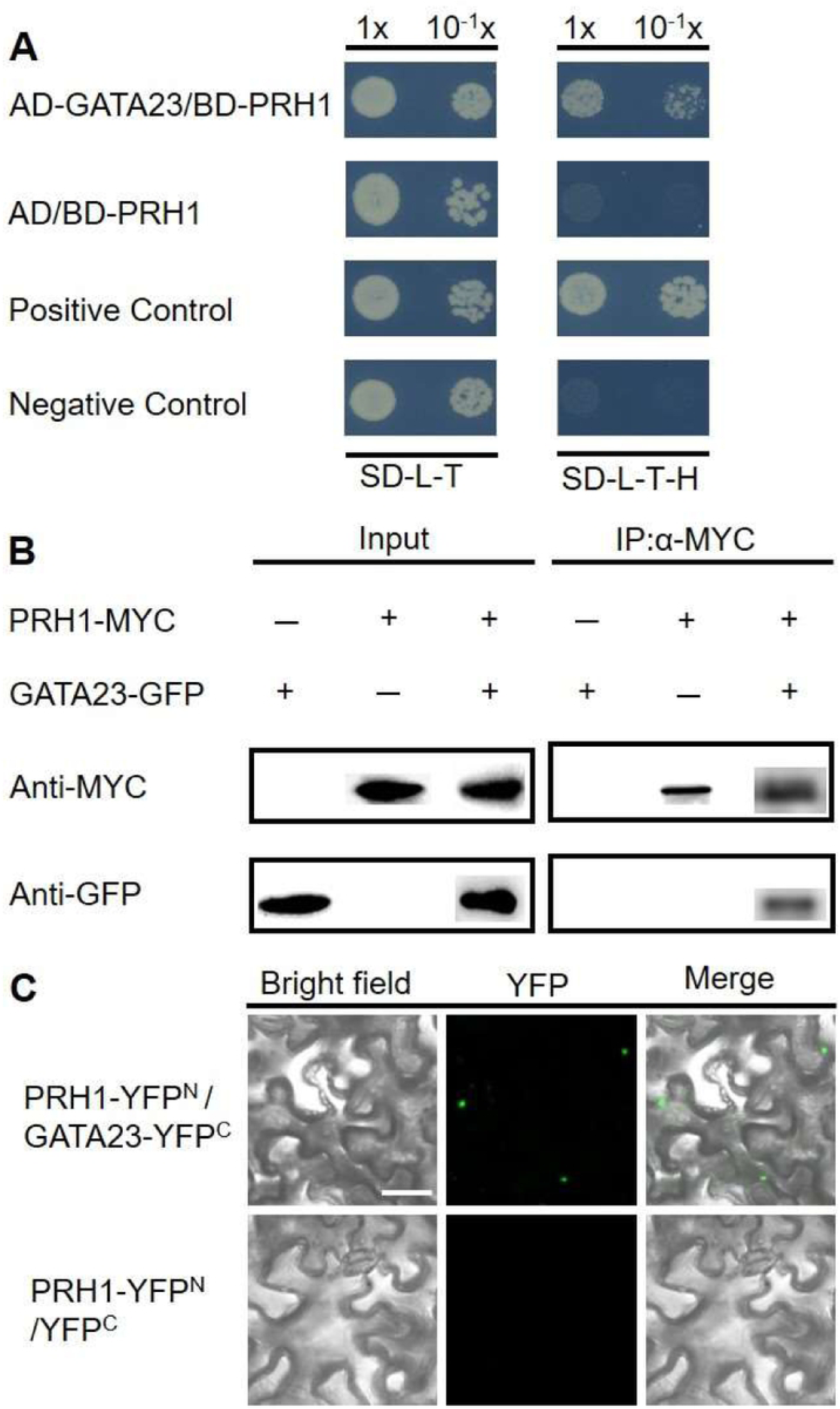
PRH1 physically interacts with GATA23. (A) PRH1 interacts with GATA23 in a yeast two hybrid assay. Yeasts were grown on both Synthetic Complete-Leu-Trp (SD-L-T) and Synthetic Complete-Leu-Trp-His (SD-L-T-H) medium. The interaction of pGADT7-GATA23 (AD-GATA23) with pGBKT7-PRH1 (BD-PRH1) was assayed by contrasting the growth of the positive and negative controls on SD-L-T-H. (B) The interaction of PRH1 with GATA23 was assayed by Co-IP. The protoplasts transiently expressing *PRH1-MYC* and/or *GATA23-GFP* were prepared from *A. thaliana* leaves, the tagged proteins were immunoprecipitated using an agarose-conjugated anti-MYC matrix, and the blots were probed with anti-GFP and/or anti-MYC antibody. (C) Bimolecular fluorescence complementation (BiFC) was used to assay the interaction of PRH1 with GATA23. *PRH1* was fused with the N-terminal fragment of *YFP* (*YFP^N^*) and *GATA23* with the C-terminal fragment of *YFP* (*YFP^C^*); the transgenes were infiltrated into *N. benthamiana* and the leaves were assayed for YFP fluorescence. Empty vectors were used as the negative control. Scale bar: 40 μm.

According to this study together with the previous reports, a proposed model was given in Fig 8. PRH1 is involved in ARF7-LBD dependent auxin signaling pathway regulating lateral root development. PRH1 acts as a downstream gene of both ARF7 and LBDs by the direct transcriptional regulation [14]. Nuclear localized PRH1 can interact with another transcription factor GATA23, known to act downstream of ARF7 and the LBDs, to mediate LR formation [8, 20].

**Fig 8.**
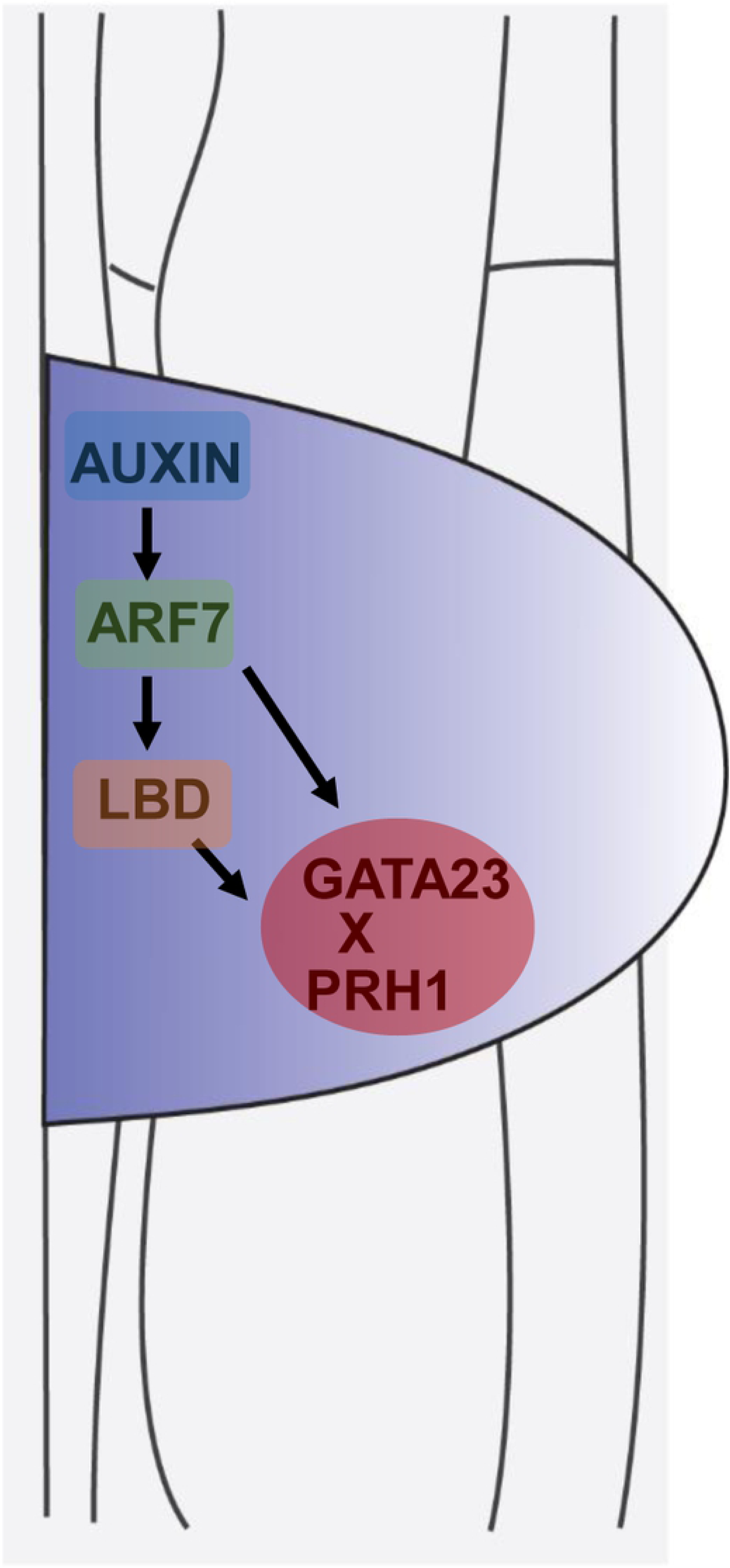
A working model of the regulation of LR development by PRH1. PRH1 acts downstream of ARF7 and LBDs. Auxin-induced *PRH1* transcription is mediated by both ARF7 and LBDs via their binding to its promoter. The transcription factor GATA23, known to act downstream of ARF7 and the LBDs, mediates auxin signaling during LR formation and interacts with PRH1.

## Discussion

### The product of the auxin-induced gene PRH1 controls LR formation

The role of auxin in LR development has been extensively investigated. During the initiation of LR, auxin signals are transduced through the two distinct AUX/IAA-ARFs modules IAA14-ARF7/ARF19 and IAA12-ARF5 [21–25]. This process establishes LR founder cell identity, which is the initial step towards the formation of a LR. Auxin is also involved in LR emergence, since it controls the activation of *IDA*, *HAE* and *HSL2* which encode cell wall remodeling enzymes and of genes encoding various expansins in the endodermal cells which lie above the LR primordia [12, 26, 27]. Members of the *LBD* gene family encode transcription factors, characterized by their LOB domain [28], which act downstream of ARF7-mediated auxin signaling to control LR formation [13, 17, 18, 29].

Here, the *PR-1* homolog gene *PRH1* was identified as a novel target of ARF7-mediated auxin signaling. Disruption of PRH1 function compromised both the initiation and emergence of LR, as demonstrated by the reduction in the number of LR developed by the *PRH1* loss-of-function mutant (Fig 3). Although the over-expression of *PRH1* in a WT background was not associated with any LR phenotype, its over-expression in *arf7*, *lbd16*, *1bd18* or *lbd29* mutant backgrounds led to a partial rescue of the compromised LR phenotype (Figs 3 and 6). The result implied that the regulation of LR formation carried out by *PRH1* likely requires a degree of coordination with other components. The *PRH1* promoter directs its product primarily to the LR primordia (as shown by the site of GUS expression in plants expressing a *PRH1pro::GUS* transgene), which is fully consistent with the proposed role of PRH1 in LR formation. Auxin treatment strongly induced the expression of the *PRH1pro::GUS* transgene in a WT (Fig 2A and 2B), but not in an *arf7* background (Fig 2C and 2D), which implied that the auxin-induced expression of *PRH1* must depend on the presence of a functional copy of *ARF7*. LBD proteins, especially LBD29, are also involved in the induction by auxin of *PRH1*. Experimentally, it was shown that both ARF7 and LBDs were able to bind to the *PRH1* promoter (Figs 4 and 5). It will be of future interest to investigate how ARF7 coordinates with LBDs to mediate auxin signaling and therefore to control the expression of *PRH1*.

### PRH1 interacts with GATA23 to control LR formation

The homo- and heterodimerization of LBD proteins are thought to be an important determinant of LR formation [29, 30]. The AtbZIP59 transcription factor has been reported to form complexes with LBDs to direct auxin-induced callus formation and LR development [31]. All these results suggest that homodimerization and heterodimerization of signaling components are important in LR formation. Here, it was shown that GATA23, a protein which influences LR formation through its regulation of LR founder cell identity [8], interacts with PRH1 (Fig 7). According to the previous study together with our results, both GATA23 and PRH1 are auxin induced, a process which is dependent on auxin response factor ARF7 (Figs 1 and 2) [8]. Whether the interaction between GATA23 and PRH1 affects the stability and/or the activity of either (or both) of these proteins remains an issue to be explored.

## Materials and Methods

### Plant materials and growing conditions

*A. thaliana* ecotype Col-0 was used as the wild type (WT), and all mutants and transgenes were constructed in a Col-0 background. The mutants comprised a set of T-DNA insert lines obtained from the Arabidopsis Biological Resource Center (ABRC, Columbus, OH, USA), namely *prh1-1* (GK_626H08), *prh1-2* (Salk_014249), *lbd16* (Salk_040739), *lbd18* (Salk_038125) [32] and *lbd29* (Salk_071133) [33], along with *arf7* (Salk_040394) [22]; the transgenic lines were *Super::LBD29-GFP* [34], *35S::MYC-LBD16*, *35S::MYC-LBD18, 35S::PRH1*, *35S::GFP-PRH1* and *PRH1pro::GUS*. Seeds were surface-sterilized by fumigation with chlorine gas, and then plated on solidified half strength Murashige and Skoog (MS) medium. All of the seeds were held for two days at 4°C and finally grown in a greenhouse delivering a 16 h photoperiod with a constant temperature of 22°C.

### Transgene constructs and the generation of transgenic lines

The 1,692 nt region upstream of the *PRH1* start cordon was used as the *PRH1* promoter, and was inserted, along with the *PRH1* coding sequence into a pENTR^TM^/SD/D-TOPO^TM^ plasmid (ThermoFisher). The construct was subsequently recombined with several Gateway destination vectors, namely PK7WG2 [35] for over-expression without any tags, pEXT06/G (BIOGLE) for over-expression with an N-GFP tag, PGWB18 for over-expression with an N-Myc tag, PBI221 for over-expression, PKGWFS7.1 [35] for promoter analysis with C-GFP/GUS, pX-nYFP (*35S::C-nYFP*)/pX-cYFP (*35S::C-cYFP*) for the bimolecular fluorescence complementation (BiFC) assay, and pGADT7 (AD)/pGBKT7 (BD) (Clontech, Mountain view, CA, USA) for the yeast hybrid assays. Transgenesis was effected using the floral dip method [36], employing transgene constructs harbored by *Agrobacterium tumefaciens* strain GV3101.

### Microscopy

Sub-cellular structures and GFP activity generated by transgenes were imaged using laser-scanning confocal microscopy. LR numbers and the GUS activity generated by transgenes were imaged by light microscopy. For the examination of LR, eight-day-old seedlings were treated following the procedure given by (Malamy and Benfey, 1997). The length of primary roots was derived from digital images using Image J software (imagej.nih.gov/ij/). The GUS staining assay followed Liu et al. [37].

### RNA isolation and quantitative real time PCR (qPCR) analysis

RNA was extracted from the roots of eight-day-old seedlings (some of which had been treated with 10 μM naphthalene acetic acid (NAA), obtained from Sigma (St. Louis, MO, USA)) using a TIANGEN Biotech (Beijing) Co. RNA simple Total RNA kit (www.tiangen.com), following the manufacturer’s protocol. A 2 μg aliquot of the RNA was used as the template for synthesizing the first cDNA strand, using a Fast Quant RT Kit (TIANGEN Biotech (Beijing) Co.). Subsequent qPCRs were processed on an MyiQ Real-time PCR Detection System (Bio-Rad, Hercules, CA, USA), using the Super Real PreMix Plus SYBR Green reagent (TIANGEN Biotech (Beijing) Co.). Each sample was represented by three biological replicates, and each biological replicate by three technical replicates. The *AtACTIN2* (*At3g18780*) sequence was used as the reference. Details of the relevant primer sequences are given in S1 Table.

### RNA-Seq analysis

RNA submitted for RNA-seq analysis was isolated from the roots of eight-day-old WT and *arf7* seedlings (both NAA-treated and non-treated samples), using a RNeasy Mini Kit (Qiagen, Hilden, Germany). The RNA-seq analysis were processed following the methods given by [38].

### Chromatin immunoprecipitation coupled to quantitative PCR (ChIP-qPCR) analysis

DNA harvested from eight-day-old seedlings of WT and transgenic lines harboring either *35S::MYC-ARF7, Super::LBD29-GFP, 35S::MYC-LBD16* or *35S::MYC-LBD18* was cross-linked in a solution containing 1% v/v formaldehyde. The ChIP procedure followed that given by [39], and employed antibodies recognizing MYC and GFP (Abcam, UK). The immune-precipitated DNA and total input DNA were analyzed by qPCR. Details of the relevant primers are given in S1 Table.

### Yeast one-hybrid (Y1H) and yeast two-hybrid (Y2H) assays

Y1H assays were carried out using the Matchmaker Gold yeast one hybrid Library Screening System (Biosciences Clontech, Palo Alto, CA, USA). The *PRH1* promoter was inserted into the *pAbAi* vector, and the bait vector (*PRH1_pro_-pAbAi*) was linearized, restricted by *Bbs*I, and introduced into the Y1H Goldeast strain along with the prey vector AD-ARF7. Transformants were grown on SD/-Leu-Ura dropout plates containing different concentrations of aureobasidin A (CAT#630499, Clontech, USA). The primers used for generating the various constructs are listed in S1Table. Y2H assays were performed as recommended in the BD Matchmaker Library Construction and Screening kit (Clontech, USA). The complete coding sequence of *PRH1* was fused to the GAL4 DNA binding domain (BD) (pGBKT7), and that of *GATA23* to the GAL4 activation domain (AD) (pGADT7), and the two constructs were introduced into yeast strain Y2HGold. Interactions were detected following the manufacturer’s protocol (Clontech, USA). The primers used for generating the various constructs are listed in S1 Table.

### Dual-luciferase transient expression assay in A. thaliana protoplasts

Protoplasts were isolated from three week old WT leaves following Yoo et al. [40]. The ligation of the *PRH1* promoter to pGreen0800-LUC was realized by enzyme digestion and ligation. The primers used for this procedure are listed in S1 Table [41]. The *PRH1pro::LUC* reporter plasmid was transferred into Arabidopsis protoplast cells with one or more effector plasmids (*ARF7, LBD16, LBD18, LBD29* – PBI221); the empty vector PBI221 was used as the negative control. The dual-luciferase assay kit (Promega, USA) allows for the quantification of both firefly and renilla luciferase activity. Signals were detected with a Synergy 2 multimode micro-plate (Centro LB960, Berthold, Germany).

### Bimolecular fluorescence complementation (BiFC) and co-immunoprecipitation (Co-IP) assays

For the BiFC assays, *Ag. tumefaciens* (strain GV3101) transformants carrying either *PRH1-YFP^N^* or *GATA23-YFP^C^* were co-infiltrated into the leaves of *N. benthamiana*, or with an empty vector. After incubation, the plants were kept in the dark for 24 h, then moved to a lit greenhouse for 72 h. The leaves were monitored by laser scanning microscopy in order to detect fluorescent signals generated by YFP. For the Co-IP assay the target genes were constructed to the over-expression vectors [42], and the plasmids of *35S::PRH1-GFP* and *35S::GATA23-MYC* were co-transformed into *A. thaliana* protoplasts using the method described above, then the protoplasts were held for 12-16 hours in darkness with the temperature of 22°C. The protoplasts were collected and the subsequent work was operated following the protocol for immunoprecipitation of Myc-fusion proteins using Myc-Trap®_MA (Chromotek, Germany). Both the input and the IP samples were separated by SDS-polyacrylamide gel electrophoresis, then transferred on to a membrane and probed with the relevant antibody.

### Statistical analysis

Tests were applied to the data to check for normality and homogeneity before attempting an analysis of the variation. The data were analyzed using routines implemented in Prism 5 software (GraphPad Software). The Students’ *t-*test was applied to compare pairs of means. Values have been presented in the form mean±SE; the significance thresholds were 0.05 (*), 0.01 (**) and 0.001 (***). Comparisons between multiple means were made using a one-way analysis of variance and Tukey’s test.

### Accession numbers

Sequence data referred to in this paper have been deposited in both the Arabidopsis Genome Initiative and the GenBank/EMBL databases under the accession numbers: *ARF7* (*At5g20730*, NP_851046), *PRH1* (*At2g19990*, NP_179589), *LBD16* (*At2g42430*, NP_565973), *LBD18* (*At2g45420*, NP_850436), *LBD29* (*At3g58190*, NP_191378), *GATA23* (*At5g26930*, NP_198045), The RNA-seq data are available in the Gene Expression Omnibus database under accession number GSE122355.

## Acknowledgments

We thank Prof. Yuxin Hu and Prof. Eva Benkova for sharing published materials.

## Supplemental data

**S1 Fig. Transcriptomic analysis of WT and *arf7* roots before and after auxin treatment.**

(A) Numbers of DEGs in *arf7* roots compared to WT with auxin treatment or not. (B) Selected candidate genes from both the up-regulated DEGs in WT by auxin and down-regulated genes in *arf7* (the bigger purple circle), and the 8 targets involved in lateral root development further refined by Arabidopsis eFP Browser from the 23 selected candidates (the smaller yellow circle). (C) The expression pattern and change folds of 8 targets at the site of lateral roots initiation in WT and *slr-1* under the condition of auxin treatment or not.

**S2 Fig. Expression pattern of *PRH1* in intact seedlings before (left) and after (right) a 3 h exposure to 10 μM NAA.**

GUS signals appear blue. GUS: β-glucuronidase, NAA, naphthalene acetic acid. Bar: 0.5 cm.

**S3 Fig. LR density and the transcription level of *PRH1* in wild type (WT), *prh1* mutants and its over-expression lines.**

(A) The number of LR primordia per cm along the primary root. (B) LRE density per cm along the primary root. (C) LRT density per cm along the primary root in WT, *prh1-1*, *prh1-2* and plants harboring *35S::PRH1*. LRP: LR primordia, LRP: LR primordia, LRE: emerged LRs, LRT: total LRs including the LRP and LRE. Values shown as mean±SE, three biological replicates in the experiment, 20 plant seedlings for each repeat. *: means differ significantly (*P*<0.05) from the WT control. (D) *PRH1* transcription level in the roots of WT, *prh1* mutants and plants over-expressing *PRH1*. Data represent means±SE, three biological replicates in the experiment. **, ***: means differ significantly (P<0.01, P<0.001) from the WT control.

**S4 Fig. The transcription of *PRH1* is regulated by LBD16, LBD18 and LBD29.**

(A) Transcription level of *PRH1* in the roots of WT, *lbd16*, *lbd18* and *lbd29* seedlings exposed to 10 μM auxin for 4 hours (orange) or not treated (brown) before extracting total RNA. Different letters atop the columns indicate significant (*P*<0.05) differences in abundance. (B) Transcription level of *PRH1* is enhanced in each of the *LBD16*, *18* and *29* over-expression lines. Data represent means±SE, three biological replicates in the experiment **, ***: means differ significantly (P<0.01, P<0.001) from the WT control.

**S5 Fig. The interaction between LBDs and *PRH1* promoter in yeast one-hybrid.**

(A) Structure of the *PRH1* promoter, showing the putative binding motifs of LBD18 (red square) and LBD29 (yellow square). (B) Yeast one-hybrid binding assay containing the interaction between LBD16, LBD18, LBD29 and *PRH1* promoter.

**S6 Fig. PRH1 is deposited in the nuclei of *N. benthamian*a leaf cells.** A *35S::GFP-PRH1* transgene was infiltrated into *N. benthamian*a leaves, and epidermal cells were scanned for the expression of GFP. GFP: green fluorescent protein. Scale bar: 20 μm.

**S7 Fig. The transcription level of *GATA23* is repressed in the roots of the mutants *arf7*, *lbd16*, *lbd18* and *lbd29*.**

Data represent means±SE, three biological replicates in the experiment. *, **: means differ significantly (P<0.05, P<0.01) from the WT control.

**S1 Table. Primers used in this study.**

